# Cardiac patch treatment alleviates ischemic cardiomyopathy correlated with reverting Piezo1/2 expression by unloading left ventricular myocardium

**DOI:** 10.1101/2022.05.08.490841

**Authors:** Yang Zhu, Yuwen Lu, Sibo Jiang, Ting Shen, Chengbin He, Yun Gao, Liyin Shen, Qiao Jin, Yuting Zhao, Hongjie Hu, Jun Ling, Jin He, Lenan Zhuang

**Affiliations:** MOE Key Laboratory of Macromolecular Synthesis and Functionalization, Department of Polymer Science and Engineering, Zhejiang University, Hangzhou 310027, China; College of Animal Sciences, Zhejiang University, Hangzhou 310058, China; Department of Radiology, Sir Run Run Shaw Hospital, Zhejiang University School of Medicine, Hangzhou, Zhejiang, 310016, China; Key Laboratory of Cardiovascular Intervention and Regenerative Medicine of Zhejiang Province, Department of Cardiology, Sir Run Run Shaw Hospital, College of Medicine, Zhejiang University, Hangzhou, 310016, China

## Abstract

Pathologically elevated mechanical load promotes the adverse remodeling of left ventricle (LV) post myocardial infarction, which results in the progression from ischemic cardiomyopathy to heart failure. Cardiac patches could attenuate adverse LV remodeling by providing mechanical support to infarcted myocardium and border zone tissue. However, the mechanism of the translation from mechanical effects to favorable therapeutic outcome is still not clear. By transcriptome analysis, we found that the myocardial transcription levels of mechanosensitive ion channel proteins Piezo1 and Piezo2 significantly increased in patients with ischemic cardiomyopathy. In vitro tensile tests with local tissue information revealed a significant decrease in local strain and mechanical load in rat infarct. Cardiac function and geometry were preserved compared to non-treated control. Further, in LV myocardium of the patch-treated group, MI induced expression levels of Piezo1/2 were significantly reverted to the similar levels of the Sham group, indicating that cardiac patch beneficial effects were correlated with suppressing mechanosensitive genes, particularly Piezo1/2. These findings demonstrated the potential of cardiac patches in treating ICM patients with remodeling risks, and could provide guidance for improvement in next generation of patch devices.

## Introduction

Myocardial infarction is one of the major threats to human health. Loss of cardiomyocytes due to lack of oxygen supply leaves a necrotic, passive region in ischemic left ventricle. To compensate for the lost function of the necrotic myocardium, the border zone and remote myocardium output greater contraction force, imposing higher wall stress in the injured myocardium[1, 2]. Together with post-MI chronic-inflammation and fibrosis, the passive and cyclic stretching under increased wall stress induces pathologically remodeling of the necrotic myocardium and border zone tissue, featured with cardiac wall thinning and LV dilation[3, 4]. As described by Laplace’s law, the increased wall stress, decreased wall thickness and volumetric increase of LV form a vicious cycle which self-propels the progression of remodeling, resulting in LV aneurysm and heart failure[5].

Epicardial implantation of mechanically strong, elastic patches covering the infarct and border zone myocardium could break the vicious cycle of adverse LV remodeling by lowering cardiac wall stress, as the patch materials share the mechanical load, increase wall thickness, and limit diastolic movements of LV wall[6]. Maintenance of LV function and geometry by patch devices has been reported in a great number of large and small animal studies[7, 8]. However, the mechanical effect based theory could not fully explain the observed therapeutic effects lacking connection by signaling pathways. Particularly, the mechano-sensors need to be identified, as the initiators on the cascade of reactions translating the mechanical influences of patches to corresponding biological changes. As a mechanosensitive ion channel protein, Piezo1 (Piezo type mechanosensitive ion channel component 1) serves as an important cardiac mechanotransducer maintaining normal heart function. Recently, Jiang et al. reported that Piezo1 expression elevates in hypertrophic cardiomyopathy patients, which demonstrated the important role Piezo1 plays in pathological cardiac remodeling[9].

We hypothesize that the expression levels of Piezo1/2 significantly increase in infarcted left ventricles as a result of pathologically elevated myocardial mechanical load, and mechanical support from cardiac patches could decrease local stress and strain, hence mediate Piezo1/2 levels in the injured myocardium. Integrated analysis of the transcriptome of cardiac biopsy from ischemic cardiomyopathy (ICM) patients and healthy control revealed elevated expression of Piezo1/2 in the remodeled LV. The mechanical effects of an elastic polyester patch on infarcted myocardium were directly measured on ex vivo rat hearts in stretch tests, supported by dot array tracking based local strain analysis. Meanwhile, the patch treatment significantly preserved the heart function and alleviated fibrosis. Further, genome wide transcriptional analysis by RNA sequencing deciphered Piezo1/2 expression levels increased in MI animals could be reverted by patch treatment. Together, our data demonstrated that ischemia induced remodeling of LV increased the expression of Piezo1/2 and cardiomyopathy genes, while the cardiac patch treatment could protect against the pathogenesis by unloading the stretch and reverting the expression of Piezo1/2 in left ventricular myocardium.

## Materials and methods

### GEO data analysis

The online human datasets, which comprised human heart tissue RNA-seq results from 29 non-failing (Ctrl) donors, and 31 ischemic cardiomyopathy patients (ICM), were obtained from Gene Expression Omnibus (GEO, Accession No.: GSE116250, GSE120852 and GSE46224). The downloaded sra-formatted files were first converted to fastq format using fastq-dump command. Then, these files were processed using the same protocol to the rat’s data described belowFastp was used to generate clean data by removing low-quality reads, poly-N reads, and adapter sequences. Then, the clean data of rat and human were aligned to their respective reference genomes (rat genome version 6.0.104, human genome version GRCh38) using Hisat2 tool[10]. Subsequently, the samtools[11] was employed to transform the aligned data to sorted bam files, which were counted by featureCounts function from Rsubread package.

For humans, the raw counting matrices were processed and reclassified to have four groups: Ctrl, ICM for rats, batch effects were included in the normalization function. DESeq2[12] was utilized to normalize these data, and the differentially expressed genes (DEGs) were defined as those genes which had adjusted *p* value < 0.05. The expression levels of PIEZO1 and PIEZO2 genes of human and rat extracted from the normalized data were analyzed by one-way ANOVA and Tukey’s HSD post hoc. Then, principal component analysis (PCA) and heatmap were depicted after removing batch effect by removeBatchEffect of limma package [13] and stabilizing variances by vst function[12].

Gene ontology (GO) analyses of differentially expressed genes were carried out by the Database for Annotation, Visualization and Integrated Discovery (DAVID). To further explore the functional annotations of the DEGs, gene set enrichment analysis (GSEA) was conducted using the clusterProfiler package, and the gene sets related to mechanical stimulation and cardiac contraction were included in the analysis.

### Synthesis of multiblock copolymers

Tetrahydrofuran (THF, Sinopharm Chemical Reagent, China) was refluxed over potassium to remove the moisture. Propylene oxide (PO, 98%, Energy Chemical, China) and ε-caprolactone (CL, 99%, Acros, China) were distilled over CaH_2_ (Sinopharm Chemical Reagent, China) and stored in argon. Lu(OTf)_3_ was synthesized by hydrothermal reaction of Lu_2_O_3_ (>99.99%, Founde Star Science & Technology, China) and triflic acid (99%, Energy Chemical, China) as reported[14] and dried in vacuum at 200 °C for at least 40 h. PCL-*b*-p(THF-*co*-CL)]_m_ was synthesized via Janus polymerization as previously described[15]. Lu(OTf)_3_ (0.3732 g, 0.6 mmol) was added into a flame-dried ampoule and dissolved in THF (8.35 g, 116 mmol) and CL (6.84 g, 60 mmol). PO (52.2 mg, 0.9 mmol) in THF (0.29 g, 4 mmol) was then added to initiate the polymerization. The ampoule was sealed and kept at 25 °C for 14 days. The crude product was precipitated in cold methanol, filtrated and dried under vacuum.

### Characterization of the copolymer

Molecular weights and dispersities (Ð) were determined by size-exclusion chromatography (SEC) on Waters-150C equipment with Waters Styragel HR3 and HR4 columns and a Waters 2414 refractive index detector. THF was used as the eluent at a flow rate of 1.0 mL/min at 40 °C. Commercial polystyrene samples were used as calibration standard. Nuclear magnetic resonance (NMR) spectra were recorded on a Bruker Avance DMX 400 spectrometer (^1^H: 400 MHz). CDCl_3_ was solvent and tetramethylsilane (TMS) was internal reference. Analyses was performed by differential scanning calorimetry (DSC) on a Q200 TA Instrument. Samples were heated from −40 °C to 80 °C at a heating rate of 10 °C/min under nitrogen, held for 5 min to erase the thermal history, and then cooled to −40 °C and held for another 5 min. The second heating scan from −40 °C to 80 °C at the same heating rate was recorded.

### Fabrication of the patches

[PCL-*b*-p(THF-*co*-CL)]_m_ patches were dried from 15 wt% THF solutions in Teflon molds. THF was slowly evaporated at room temperature for 48 h. After the film was removed, 8 mm patches were cut using a hole punch.

### Degradation rate of patches

[PCL-*b*-p(THF-*co*-CL)]_m_ patches (8 mm diameter) were weighed (W_0_), then incubated in PBS at 37 °C, shook at 100 rpm (n = 3). Samples were rinsed 3 times with deionized water, freeze-dried for 1 day before recording the weight (W_1_) and change the PBS solution once every month. Mass remaining (%) was calculated as W_1_ / W_0_ × 100%.

### Mechanical property measurement

Mechanical properties of the patch samples before and after 3 months degradation were measured by an Instron 5540A universal testing machine. Stretching speed was 10 mm/min. Cyclic stretch test was conducted by stretching the samples to a strain of 10%, and releasing back to 0% strain for 10 cycles.

### Mechanical support by the patches on infarcted myocardium ex vivo

The microneedle array seal with 9×12 microneedles was designed with Solidworks. The microneedles were cone-shaped, whose bottom diameter was 0.5 mm and height was 1 mm. Space between adjacent microneedles was 1 mm on both X and Y directions. The seal microneedle array was 3D printed with the photosensitive resin (BMF Precision Tech, China).

All animal experiments were approved by the Guidelines of ZJU Animal Protection and Use Committee (20150119-013). Male Sprague Dawley rats (8-10 weeks, 200-250 g) were anesthetized with 1% pentobarbital. Left ventricular infarction was created by ligating the left anterior descending (LAD) coronary artery with a 6-0 filament. After 30 minutes of ligation, the left ventricular myocardium was excised, unfolded circumferentially and marked with the seal. Distal ends of the left ventricular myocardium were fixed to hooks on the upper and lower clamps of a universal mechanical testing machine (Instron, USA). The left ventricular myocardium was stretched at a rate of 1 mm/s to reach 10% strain. Subsequently, a patch was fixed to the myocardium sample with sutures, and the patched myocardium stretched and recorded using the same protocol. Myocardium samples from Sham group (myocardium punctured by needle, no ligation) was used as control group.

The microneedle array seal was dipped in the fluorescent dye solution (Xiucai Chemicals, China) and transferred a dot array to the surface of myocardium samples. The samples were stretched under ultraviolet light, and the movement of the fluorescent dots were recorded by a fixed camera. Pictures before and after stretching were taken from captured from corresponding videos. Distances between the dots on both X and Y directions before and after stretching were measured, and used to calculate local strains in circumferential and longitudinal directions. The strain map was visualized with color levels, in which the color depth reflected the local strain level. The effects of the patch on stress, strain on circumferential and longitudinal directions, mechanical load on the infarcted myocardium, as well as the complex modulus of the myocardium/patch composite at 10% total stretching were calculated.

### Rat myocardial infarction model and patch treatment

Male Sprague Dawley rats aged 8-10 weeks (200-240 g) were purchased from Zhejiang Academy of Medical Sciences. Myocardial infarction was created as described above. In the P (Patch) group, the patches were sutured on 4 points on the edge of the infarcts with 6-0 silk threads. Sham group, MI group (no treatment), and patch implantation with 1 stitch were used as controls. Finally, the chest was closed with 3-0 sutures. After 28 days of treatment, cardiac function was assessed using an ultrasound system (VEVO2100 ultrasound system, Visual Sonics, Canada). Left ventricular ejection fraction (LVEF), left ventricular fractional shortening (LVFS), end-diastolic volume (EDV) and end-systolic volume (ESV) were calculated.

### RNA-seq experiment

We extracted total RNA of tissues from MI sites for MI and MI treated by P groups using TRIzol (Invitrogen, USA). In the Sham group, tissues from the similar left ventricle site in the MI group were used. A total amount of 1 μg RNA per sample was used as input material for the RNA sample preparations. Sequencing libraries were generated using NEBNext UltraTM RNA Library Prep Kit for Illumina (NEB, USA) following manufacturer’s recommendations and index codes were added to attribute sequences to each sample. Briefly, mRNA was purified from total RNA using poly-T oligo-attached magnetic beads. Fragmentation was carried out using divalent cations under elevated temperature in NEBNext First Strand Synthesis Reaction Buffer (5X). First strand cDNA was synthesized using random hexamer primer and M-MuLV Reverse Transcriptase (RNase H-). Second strand cDNA synthesis was subsequently performed using DNA Polymerase I and RNase H. Remaining overhangs were converted into blunt ends via exonuclease/polymerase activities. After adenylation of 3’ ends of DNA fragments, NEBNext Adaptor with hairpin loop structure were ligated to prepare for hybridization. In order to select cDNA fragments of preferentially 250∼300 bp in length, the library fragments were purified with AMPure XP system (Beckman Coulter, Beverly, USA). Then 3 μl USER Enzyme (NEB, USA) was used with size-selected, adaptor-ligated cDNA at 37°C for 15 m in followed by 5 min at 95 °C before PCR. Then PCR was performed with Phusion High -Fidelity DNA polymerase, Universal PCR primers and Index (X) Primer. At last, PCR products were purified (AMPure XP system) and library quality was assessed on the Agilent Bioanalyzer 2100 system. All the sequence data have been deposited in NCBI’s Gene Expression Omnibus (GEO, http://www.ncbi.nlm.nih.gov/geo) and are accessible through GEO series accession number GSE202228.

### RNA Extraction, Reverse Transcription, and quantitative Real-Time PCR

Total RNA was isolated from rat heart tissues using an TRIZOL method. cDNA was obtained using a EZB Reverse Transcription System (EZBioscience, USA) and analyzed by quantitative Real-Time PCR (qRT-PCR) using SYBR Green (TSINGKE, China). The data were normalized to expression of 18S. The primer sequences are shown in **Table 1**.

### Statistics

Graphpad Prism was used for statistical analysis. Results are expressed as means ± SEM (standard error of the mean). The statistical differences were tested with analyzed using one-way analysis of variance (ANOVA), when there were three groups or more; or the data were analyzed by unpaired t-test. Type I error was set at 0.05. \**p* < 0.05; \*\*\*\**p* < 0.0001, two-tailed unpaired Student’s t-test.

## Results

### Expression of Piezo1/2 significantly increased in ICM patients

To discover whether the mechanosensitive genes were changed in ischemic cardiomyopathy (ICM) patients, we combined 3 ICM datasets (GSE116250, GSE120852 and GSE46224) with batch effects removed (**Figure 1A**). The principal component analysis (PCA) showed that ICM significantly changed the transcriptome compared with the Ctrl (**Figure 1B**). Meanwhile, volcano plot visualized the differentially expressed genes (DEGs) including up-regulated 1035 genes and down-regulated 850 genes by comparing the ICM group with the Ctrl group (**Figure 1C**). The expression levels of genes related to mechanical stimulus represented by PIEZO1 and PIEZO2 genes significantly increased in the myocardial tissues of patients with ICM, whereas the expression levels of genes involved in the regulation of cardiac muscle cell contraction pathway represented by CACNA1C and ACTC1 genes decreased in ICM patients (**Figure 1D, E**).

**Figure 1.**
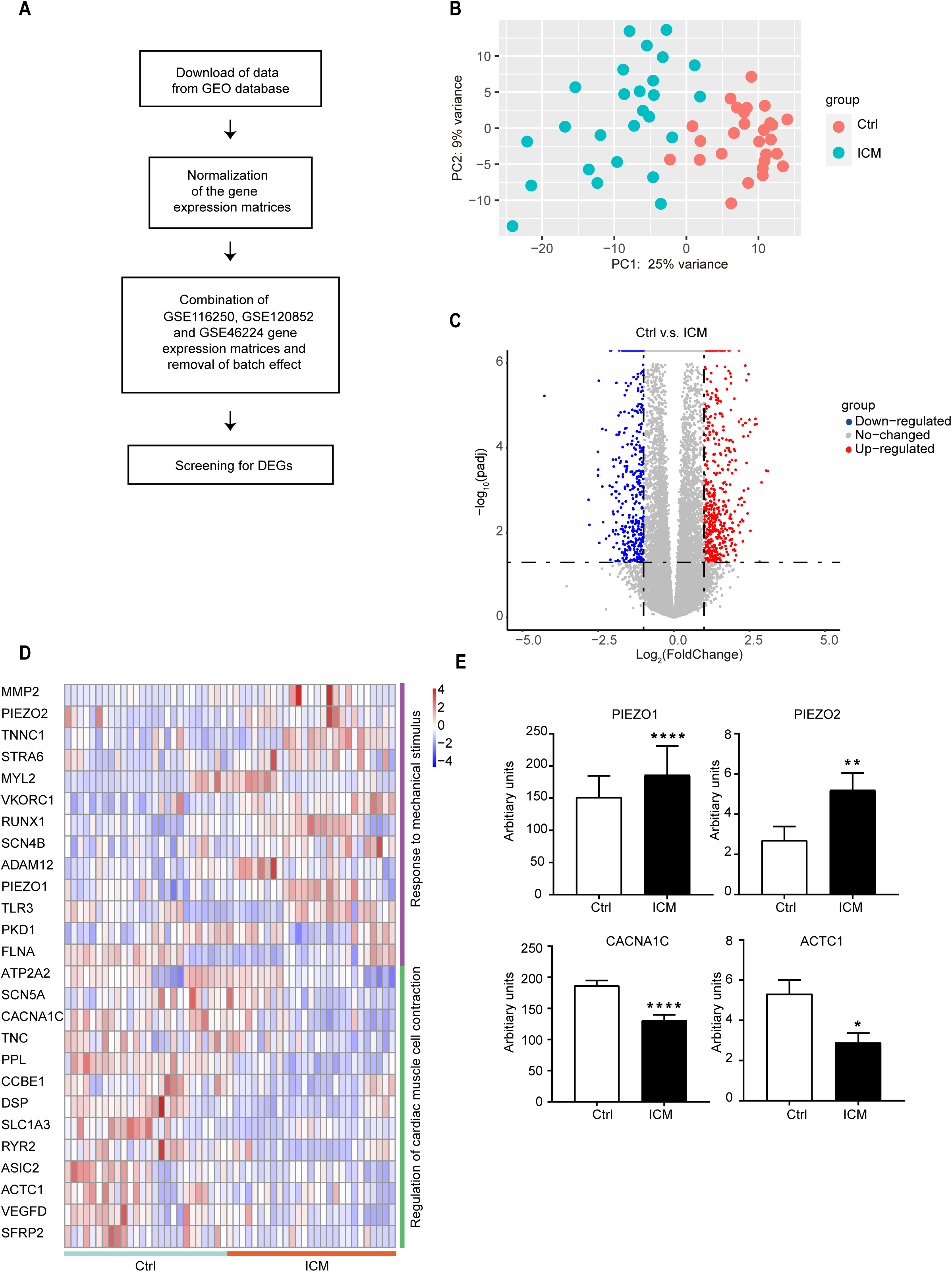
Significantly altered gene expression profiles in the ICM patients. (A) Workflow of the study design. (B) Principal component analysis (PCA) of merged RNA-seq data indicated the separation of control (Ctrl) and ICM individuals. (C) Volcano plot of all expressed genes from merged RNA-seq data. Horizontal dash-dot line represents the threshold for adjusted *p* value = 0.05, while vertical dash-dot lines show the Log_2_ (FoldChange) = 1. (D) Heatmap presentation of the mechanical stimulus and cardiac muscle contraction related genes. (E) Expression of PIEZO1, PIEZO2, CACNA1C, ACTC1 in the Ctrl and ICM patients. Means ± SEM. of the original data are presented. \**p* < 0.05; \*\**p* < 0.01; \*\*\*\**p* < 0.0001.

### [PCL-*b*-p(THF-*co*-CL)]_m_ patches were elastic and stable

In **Scheme 1**, [PCL-*b*-p(THF-*co*-CL)]_m_ was synthesized via Janus polymerization of ε-caprolactone (CL) and tetrahydrofuran (THF) with lutetium triflate/propylene oxide (Lu(OTf)_3_/PO) as the catalytic system. Its structure was confirmed by ^1^H NMR spectra with well assignment of each peak (**Figure 2C**). PCL is a semicrytalline polymer with a T_m_ of 59-64 °C and T_g_ of −60 °C[16]. During the second heating scan in DSC test, the melting temperature (T_m_) was observed at 57 °C. (**Figure 2A**). The *M*_n_ of the obtained multiblock copolymers ([PCL-*b*-p(THF-*co*-CL)]_m_) is 203.0 kDa and the PDI is 1.8 (**Figure 2B**).

**Figure 2.**
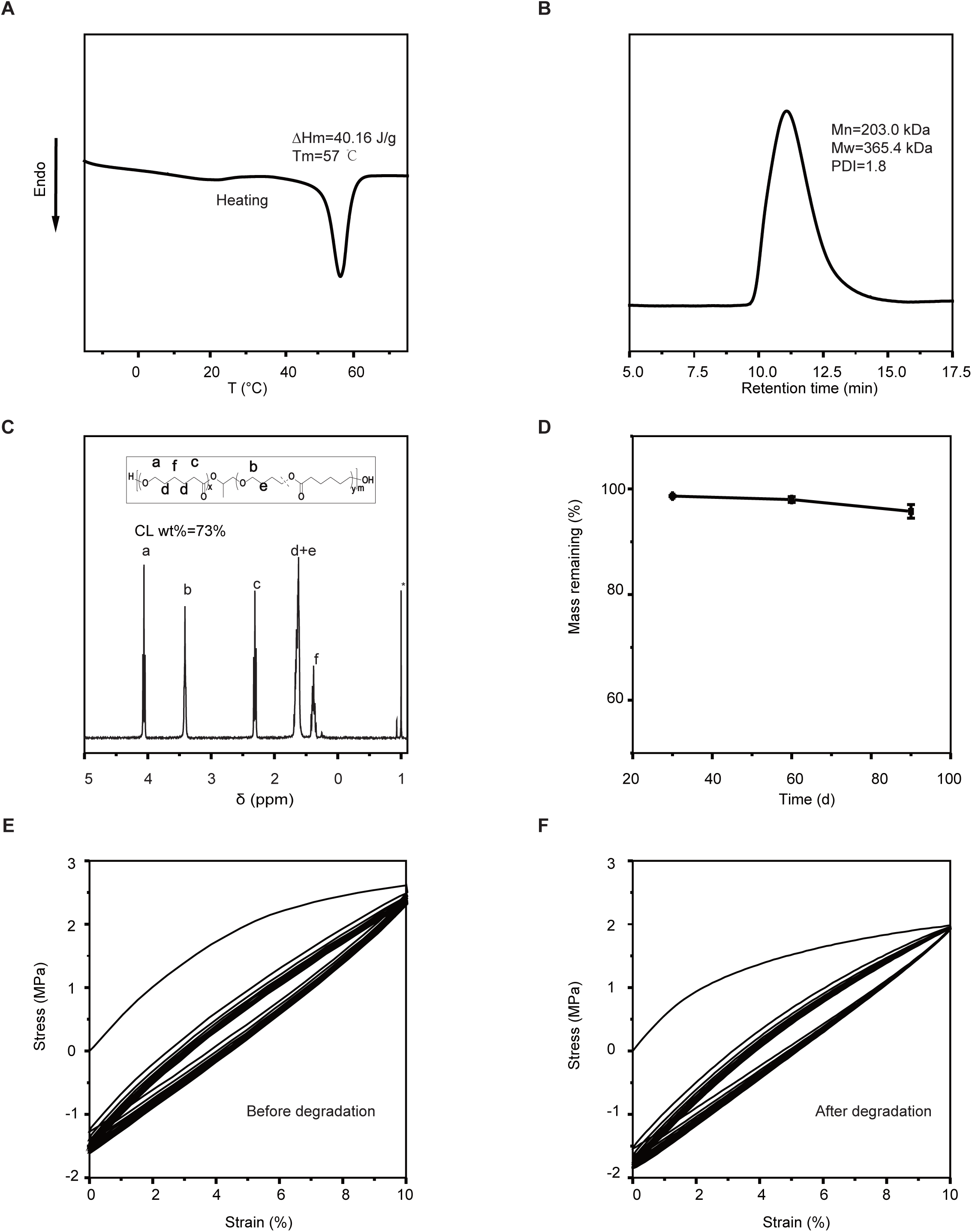
Characterizations of PCL-*b*-p(THF-*co*-CL)]_m_. (A) DSC curve of PCL-*b*-p(THF-*co*-CL)]_m_. (B) SEC traces of PCL-*b*-p(THF-*co*-CL)]_m_ for molecular weight measurement. (C) ^1^H NMR spectra of PCL-*b*-p(THF-*co*-CL)]_m_. (D) Mass remaining (W_t_/W_0_*100%) of PCL-*b*-p(THF-*co*-CL)]_m_ after different degradation time. Cyclic stretch results of cardiac patches at 10% elongation before (E) and after (F) degradation.

In PBS, the ester bonds in the patch were randomly hydrolyzed and cleaved. After 90 d degradation, the mass remaining of [PCL-*b*-p(THF-*co*-CL)]_m_ patch was 95.8%, showing a small weight loss (**Figure 2D**). To mimic the cyclic mechanical strains on the heart, the patches were cyclically stretched at 10% strain (**Figure 2E, F**). None of the patches or the stitch points failed, and no irreversible deformation after 90 d degradation was observed. Compared to fresh patches, 73% original stress was needed to obtain the same 10% strain. The exhibited stability of [PCL-*b*-p(THF-*co*-CL)]_m_ patch is expected to maintain the mechanical support in vivo.

### Cardiac patches provided mechanical support and decreased the local strains in the infarct tissue

When stretched at 10% strain along the circumferential direction of the myocardium, the healthy tissue in the Sham group had 9.3% strain in the circumferential (Y) direction and −9.0% strain in the longitudinal (X) direction (**Figure 3A, B**). In MI group, the infarct area had greater strains in the circumferential (Y) direction (12.9%) and in the longitudinal (X) direction (−12.5%) (**Figure 3A, B**) compared to the healthy region on the same sample and the Sham control. The cardiac patches reduced circumferential strain to 7.6% and the longitudinal strain to −7.8% in the infarct area (**Figure 3A, B**), smaller compared to the Sham group. As calculated from strain, total load on the myocardium samples, the stress and mechanical load in the infarct were reduced, and the modulus was increased as a result of patch implantation. Compared to the same myocardium before suturing the patches, cardiac wall stress in circumferential direction was reduced from 11.3 kPa to 8.8 kPa (**Figure 3D**), mechanical load in the myocardium was reduced from 0.28 N to 0.22 N (**Figure 3C**), and the complex modulus was increased from 69.5 kPa to 93.8 kPa (**Figure 3E**). The reduction in local strains after patch implantation was smaller in the remote area, compared to the infarct and border zone tissue covered by the patch, showing that the mechanical support effects mainly concentrated in the patched area.

**Figure 3.**
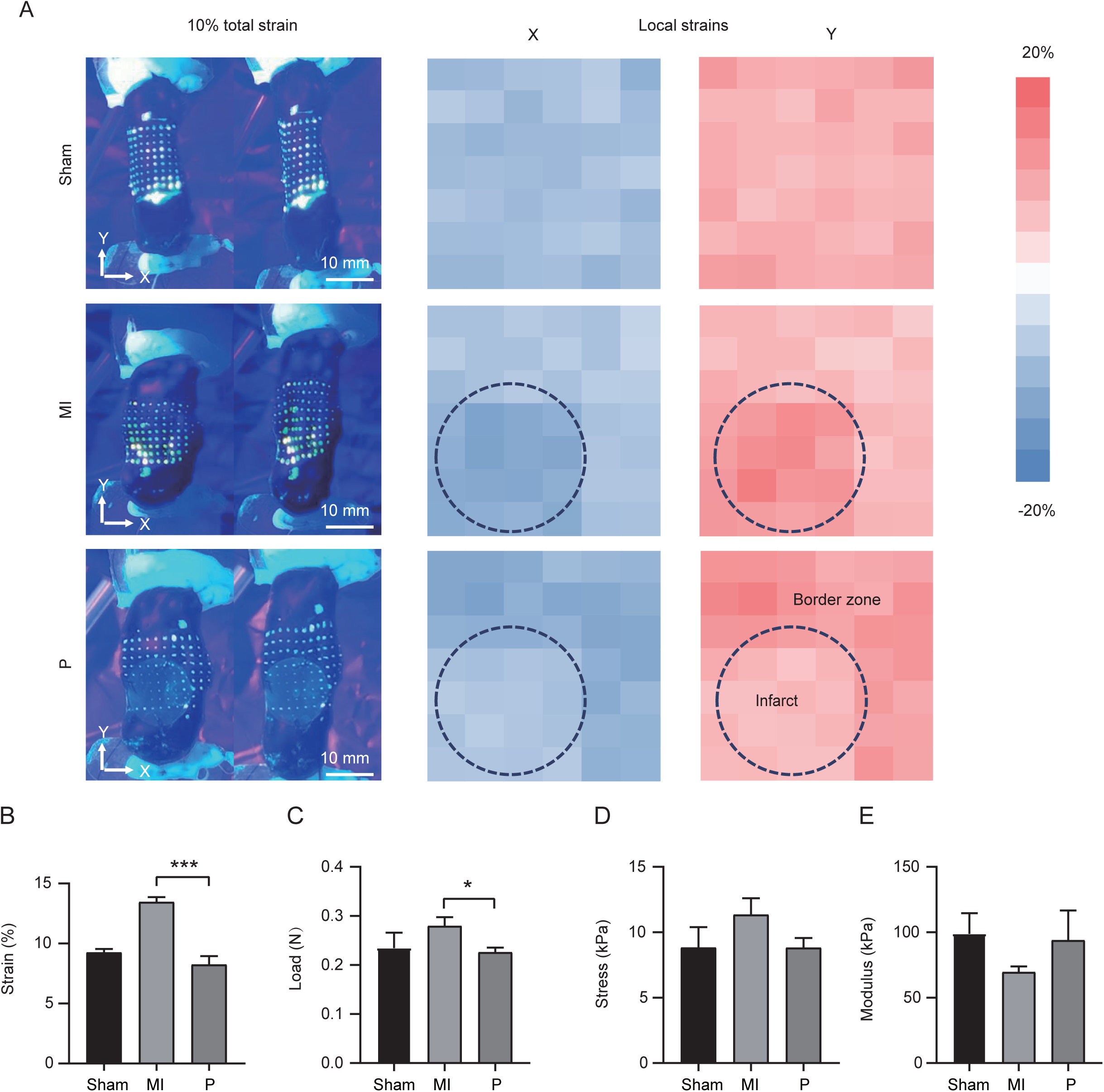
Influence of patch on local myocardial strain and mechanical load ex vivo. (A) Images of excised myocardium before and after 10% uniaxial stretch along the circumferential direction (left column). The right column shows the local strain distribution in X (longitudinal, blue) and Y (circumferential, red) directions on stretched (10% total strain) rat myocardium (Sham, MI, and P). (B) Strain, (C) load (D) stress, and (E) composite modulus of infarct region. \**p* < 0.05, \*\**p* < 0.01 and \*\*\**p* < 0.001 versus MI group.

### Cardiac patches preserved cardiac functions and attenuated adverse remodeling of LV

Severe fibrosis (25.3%) was observed in the MI group (**Figure 4G**), as shown by Masson’s trichrome staining. After 28 days of patch treatment, fibrosis area was 21.4% (1 stitch) and 18.9% (3 or 4 stitches), significantly smaller compared to MI control (**Figure 4F, G**). Echocardiography images were applied to observe the restoration of patches on cardiac functions **(Figure 4A**). Left ventricular ejection fraction (LVEF) and fraction shortening (LVFS) of rats from MI group were 38.9% and 16.5%, respectively. Patch implantation with 3 or 4 stitches maintained the LVEF and LVFS at 53.8% and 24.4%, respectively, significantly higher compared to MI control and 1 stitch patch group with the LVEF and LVFS at 46.7% and 21.3% (**Figure 4B, C**). In addition, the patch alleviated LV dilation. Hearts treated by 3 or 4 stitches patches retained significantly smaller end-diastolic volume (1.0 μl, EDV) and end-systolic volume (0.5 μl, ESV) compared to MI control (1.5 μl, 0.9 μl) and 1 stitch patch group (1.3 μl, 0.6 μl) (**Figure 4D, E**).

**Figure 4.**
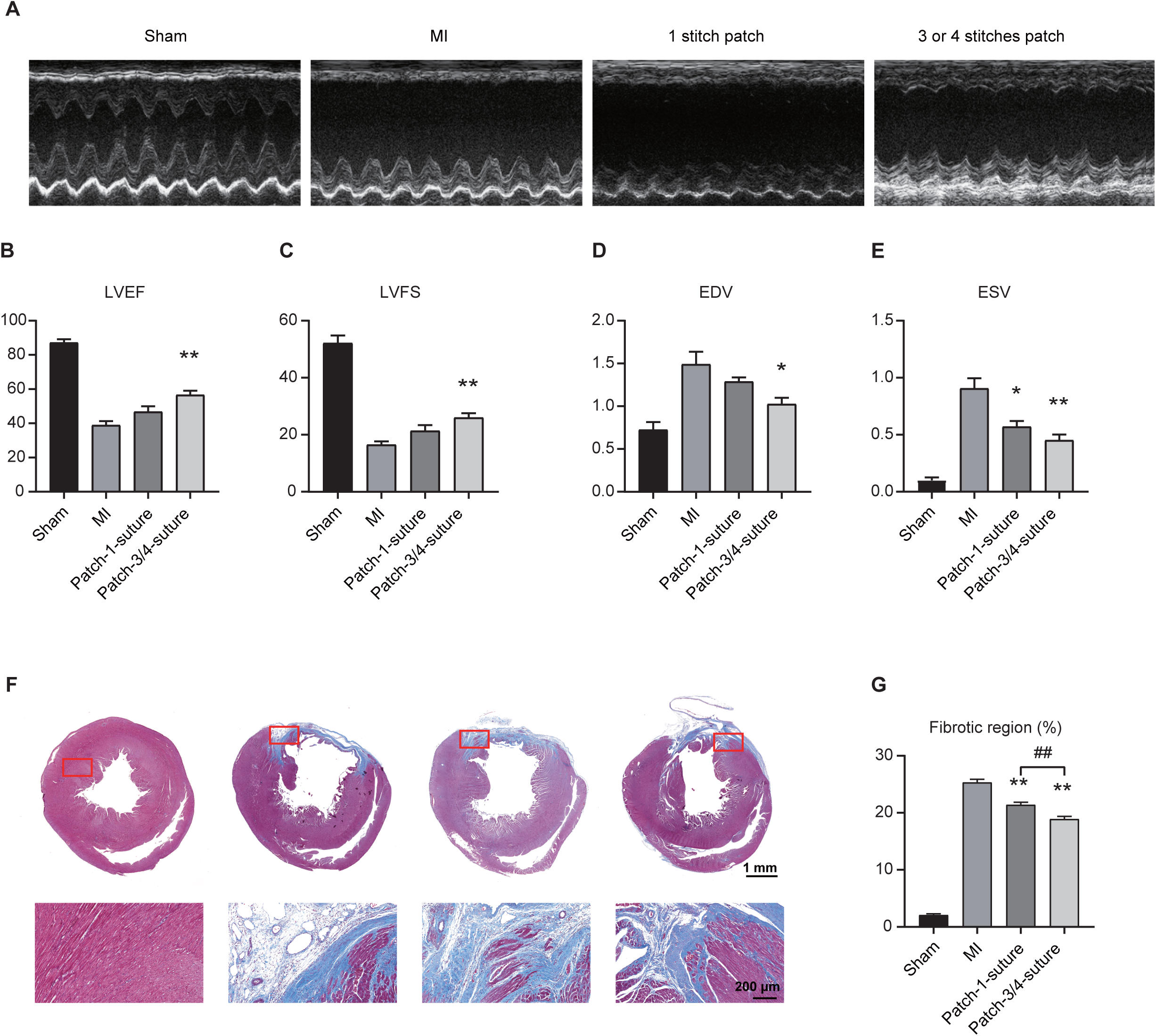
Improvement in cardiac functions of MI rats by patch treatment. (A) Representative M-mode echocardiography images in different groups 28 d post-surgery. Echocardiography analysis of (B) left ventricular ejection fraction (LVEF), (C) left ventricular fractional shortening (LVFS), (D) end-diastolic volume (EDV), and (E) end-systolic volume (ESV) 28 days after surgery. (F) Masson’s trichrome staining of the rat hearts (upper row) and local microscopic images of the border zone myocardium (red boxes, lower row) 28 days after surgery, and (G) Ratio of fibrotic region in left ventricle. \**p* < 0.05, \*\**p* < 0.01 versus MI group, ^##^*p* < 0.01 comparison as indicated in figure.

### Transcriptome analysis indicates the patch treatment preserved cardiac function after MI via reverting Piezo1/2 expression

To determine the therapeutic efficacy of the cardiac patch at the molecular level, RNA-seq was used to identify different expression genes among the three groups. Principal component analysis showed a separation in transcriptomes among the three groups (**Figure 5A**). Consistently, the correlation matrix and clustering revealed transcriptome features of P and Sham groups were clustered together, separated with the MI group, which could reflect the patch treatment changed the transcriptome of the infarct myocardium to be similar with the normal control (**Figure 5B**). Compared to the Sham group, there were 2494 upregulated genes and 882 downregulated genes after MI operation (MI v.s Sham) (**Figure 5C**). Patch treatment resulted in the up-regulation of 392 genes and the down-regulation of 627 genes (P v.s MI) (**Figure 5D**). In order to further explore the changes after treated by patch, we analyzed a variety of gene sets by the gene set enrichment analysis (GSEA). Consistent with our finding in **Figure 1**, expression of genes in response to mechanical stimulus, including *Piezo1/2*, were significantly upregulated in MI compared with Sham (**Figure 5E and H**). Particularly, cardiac patch treatment reverted these elevated gene expressions in the infarct regions in the P group (**Figure 5H**). Simultaneously, expression of genes in regulation of cardiac muscle cell contraction were significantly decreased in MI compared with Sham (**Figure 5F and I**). Nevertheless, patch treated infarct myocardium restored those cardiomyocyte contraction gene expression while preserving the heart function **(Figure 5G and I**). The RNA-seq derived expression values of the representative genes, *Piezo1/2* for the response to mechanical stimulus; *Atp1a1* for the cardiac muscle cell contraction, were shown in **Figure 5J**. Similar results were confirmed by qRT-PCR with total RNA extracted from the same samples used in the RNA-seq experiment (**Figure 5K**).

**Figure 5.**
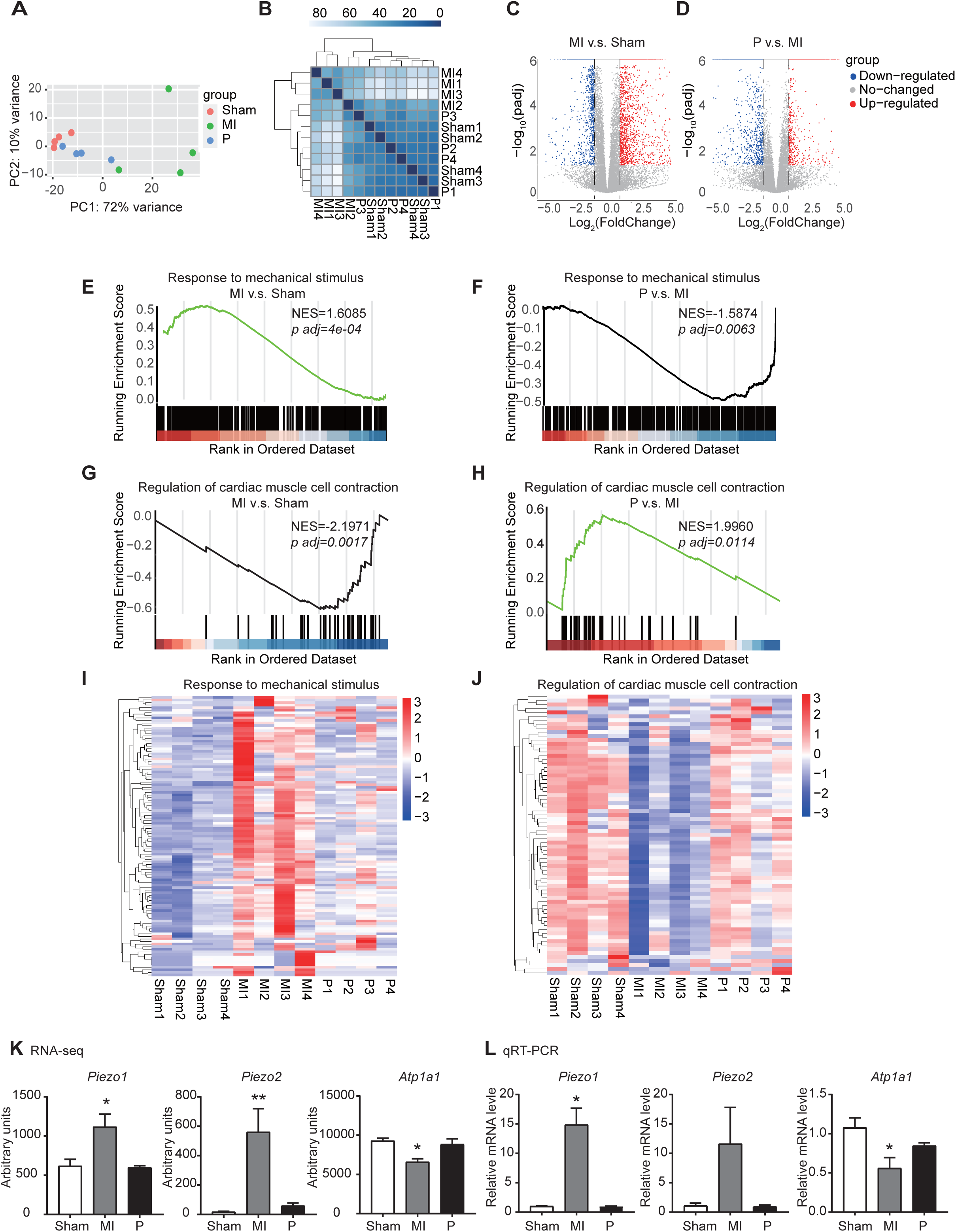
Transcriptome analysis indicates the patch treatment preserved cardiac function after MI via reverting Piezo1/2 expression (A) Principal component analysis (PCA) of transcriptomes of the infarct area from the Sham, MI and P (n = 4 per group). (B) Correlation matrix analysis of transcriptomes of the infarct area from the Sham, MI and P (n = 4 per group). The darker the color, the stronger the correlation is. (C-D) Volcano plot of all expressed genes from RNA-seq data between MI and Sham (C) or between P and MI (D). Horizontal dash-dot line represents the threshold for adjusted *p* value = 0.05, while vertical dash-dot lines show the value of Log_2_ (FoldChange) being 1. (E-F) GSEA analysis of response to mechanical stimulus between MI and Sham (I) Heatmap for response to mechanical stimulus. (G-H) GSEA analysis of regulation of cardiac muscle cell contraction between MI and Sham (G) or between P and MI (H). (J) Heatmap for regulation of cardiac muscle cell contraction. (K) Arbitrary expression units for *Piezo1, Piezo2, Atp1a1* between Sham, MI and P in RNA-seq assays. (L) Expression levels of *Piezo1, Piezo2, Atp1a1* between Sham, MI and P. qRT-PCR was performed with RNA extracted from the rats. \**p* < 0.05, \*\**p* < 0.01 versus Sham group.

## Discussion

Coronary artery disease (CAD) is the most common cause of ischemic cardiomyopathy (ICM) which eventually leads to heart failure [17]. Prolonged ischemia in myocardium irreversibly induces cardiac muscle damage resulting in cardiac remodeling and fibrosis, which decreased cardiac function. In mechanical respect, the ischemic myocardium is passively stretched by the contraction-relaxation cycle of the heart continuously under ICM status. Providing adequate mechanical support to the ischemic hearts by elastic cardiac patches is considered as a potentially translatable strategy for myocardial infraction treatment [18]. Piezo1/2 are mechanosensitive ion channels transducing mechanical stimuli into calcium signal in cardiomyocytes, which plays critical roles in normal heart function and disturbed in pathological conditions [19]. However, the changes in the expression of genes response to mechanical stimuli in the stretched myocardium were not clear. Jiang et al. reported the expression of Piezo1 increased in human hypertrophic cardiomyopathy hearts [9]. By re-visiting the combined expression profile datasets of ICM myocardium, our analysis revealed the expression of Piezo1/2 increased significantly in human ICM cardiac muscle, and such finding was repeated in a rodent MI model. We believe that the elevation in Piezo1/2 expression levels in ICM patients is a result of increased stress and mechanical load in ischemic myocardium. Interestingly, the patch treatment protected the heart function, and reverted the high expression of Piezo1/2 at the same time. Together, we have established the correlation between the mechanical stretch and the expression of Piezo1/2 in the context of cardiac patch treatment on ICM, provided a potential molecular basis underlying the relationship between mechanical stress and ischemic cardiomyopathy.

Attenuated post-MI pathological LV remodeling has been repeatedly achieved in studies using different elastomers. Fujimoto et al. first demonstrated that implanting an elastic polyurethane patch on infarcted LV could maintain cardiac function[18]. PCL, PLLA, and polyurethane urea with ester, caprolactone or ethylene oxide soft showed similar therapeutic effects in small animals[20-22]. Physical mixing with biomacromolecules including collagen, decellularized extracellular matrix, laminin, etc. could provide additional cell adhesiveness[23-27]. With a porcine study, Hashizume et al. demonstrated that the patch treatment concept is applicable in large animal MI models[28]. Silveira-Filho et al. showed that patch implantation in late stage MI mediated the risk of heart failure[29]. Lin et al. presented the importance of optimizing the viscoelasticity of the patch devices in order to improve the therapeutic outcome[4]. However, the key signaling factors and pathways involved in translating the mechanical effect to functional outcome has not been fully elucidated. In this study, we used a novel elastic polymer, [PCL-*b*-p(THF-*co*-CL)]_m_ as the representative polymer for the stability in its mechanical property after extended use[15]. [PCL-*b*-p(THF-*co*-CL)]_m_ degrades slower compared to PCL attributed to the inertness of THF component in hydrolytic environment. In addition, non-porous solvent casted films were selected to decrease the interference by cell-patch interaction on molecular biology analysis compared to porous patches fabricated by electrospinning, salt-leaching or thermally induced phase separation.

[PCL-*b*-p(THF-*co*-CL)]_m_ has a typical modulus of 51.4 MPa, compared to other tested elastomers[30]. Therefore, the results and conclusion presented in this study shall be applicable to patches with similar modulus and geometry. We have shown that as the strain in the patch myocardium decreased, strains myocardium in the remote area increased by about 68.2% (**Figure 3A**), which is a sign of increased mechanical load in the remote myocardium. If the patch is too stiff or too thick, it is expected that the strain in the patched myocardium would decrease to about 0, leaving all mechanical load to remote myocardium. Therefore, a highly stiff patch may result in concentrated stress in remote myocardium and associated tissue damages. On the other hand, inadequate mechanical support does not lead to cardiac function preservation. We implanted the patch with only one suturing point, the treated rats did not improve the cardiac function compared to the MI group (**Figure 4E**).

## Conclusion

This study indicated that, at the transcriptome level, ischemic cardiomyopathy decreased the expression of genes associated with cardiac contraction and increased the mechanical stimuli sensing gene expression, including Piezo1/2. By mechanically unloading the infarcted myocardium, cardiac patch treatment changes the transcriptome in molecular pathways leading to improved cardiac muscle cell contraction and reverted response to mechanical stimulus. Our results established a clear correlation underlying the mechanical stress with the expression of mechanosensitive genes Piezo1/2 in ICM patients, and demonstrated that cardiac patch treatment preserved cardiac function and geometry by reverting the pathological changes in myocardial transcriptomic levels of Piezo1/2 and related mechanosensing genes. Further studies are needed to understand how the cardiac patch treatment initiated the regulation in the mechanosensing gene transcription, and integrate the mechanical effects and biological responses including gene transcription at the microscopic cell and tissue levels.

## Acknowledgements

This study is financially supported by the National Key Research and Development Program of China (No. 2019YFE0117400 and 2021YFA0805902), National Natural Science Foundation of China (No. 81941003, 81873908 and 32000971), “Leading Goose” R&D Program of Zhejiang, Alibaba-Zhejiang University Joint Research Center of Future Digital Healthcare.

## Reference

[1] O.J. Mechanic, M. Gavin, S.A. Grossman, Acute Myocardial Infarction, StatPearls, StatPearls Publishing Copyright © 2022, StatPearls Publishing LLC., Treasure Island (FL), 2022.

[2] J.S. Burchfield, M. Xie, J.A. Hill, Pathological ventricular remodeling: mechanisms: part 1 of 2, Circulation 128(4) (2013) 388–400.

[3] L.A. Reis, L.L. Chiu, N. Feric, L. Fu, M. Radisic, Biomaterials in myocardial tissue engineering, J. Tissue Eng. Regen. Med. 10(1) (2016) 11–28.

[4] X. Lin, Y. Liu, A. Bai, H. Cai, Y. Bai, W. Jiang, H. Yang, X. Wang, L. Yang, N. Sun, H. Gao, A viscoelastic adhesive epicardial patch for treating myocardial infarction, Nature Biomedical Engineering 3(8) (2019) 632–643.

[5] W. Nadruz, Myocardial remodeling in hypertension, J. Hum. Hypertens. 29(1) (2015) 1–6.

[6] T. Chang, C. Liu, K. Lu, Y. Wu, M. Xu, Q. Yu, Z. Shen, T. Jiang, Y. Zhang, Biomaterials based cardiac patches for the treatment of myocardial infarction, Journal of Materials Science & Technology 94 (2021) 77–89.

[7] J. Tang, J. Wang, K. Huang, Y. Ye, T. Su, L. Qiao, M.T. Hensley, T.G. Caranasos, J. Zhang, Z. Gu, K. Cheng, Cardiac cell-integrated microneedle patch for treating myocardial infarction, Science advances 4(11) (2018) eaat9365.

[8] Z. Fan, Y. Wei, Z. Yin, H. Huang, X. Liao, L. Sun, B. Liu, F. Liu, Near-Infrared Light-Triggered Unfolding Microneedle Patch for Minimally Invasive Treatment of Myocardial Ischemia, ACS Appl Mater Interfaces 13(34) (2021) 40278–40289.

[9] Y. Jiang, X. Yang, J. Jiang, B. Xiao, Structural Designs and Mechanogating Mechanisms of the Mechanosensitive Piezo Channels, Trends Biochem. Sci. 46(6) (2021) 472–488.

[10] D. Kim, J.M. Paggi, C. Park, C. Bennett, S.L. Salzberg, Graph-based genome alignment and genotyping with HISAT2 and HISAT-genotype, Nat. Biotechnol. 37(8) (2019) 907–915.

[11] H. Li, B. Handsaker, A. Wysoker, T. Fennell, J. Ruan, N. Homer, G. Marth, G. Abecasis, R. Durbin, The Sequence Alignment/Map format and SAMtools, Bioinformatics 25(16) (2009) 2078–9.

[12] M.I. Love, W. Huber, S. Anders, Moderated estimation of fold change and dispersion for RNA-seq data with DESeq2, Genome Biol 15(12) (2014) 550.

[13] M.E. Ritchie, B. Phipson, D. Wu, Y. Hu, C.W. Law, W. Shi, G.K. Smyth, limma powers differential expression analyses for RNA-sequencing and microarray studies, Nucleic Acids Res 43(7) (2015) e47.

[14] L. You, T.E. Hogen-Esch, Y. Zhu, J. Ling, Z. Shen, Brønsted acid-free controlled polymerization of tetrahydrofuran catalyzed by recyclable rare earth triflates in the presence of epoxides, Polymer 53(19) (2012) 4112–4118.

[15] L. You, J. Ling, Janus Polymerization, Macromolecules 47(7) (2014) 2219–2225.

[16] M.A. Woodruff, D.W. Hutmacher, The return of a forgotten polymer—Polycaprolactone in the 21st century, Prog. Polym. Sci. 35(10) (2010) 1217–1256.

[17] A.K. Malakar, D. Choudhury, B. Halder, P. Paul, A. Uddin, S. Chakraborty, A review on coronary artery disease, its risk factors, and therapeutics, J Cell Physiol 234(10) (2019) 16812–16823.

[18] K.L. Fujimoto, K. Tobita, W.D. Merryman, J. Guan, N. Momoi, D.B. Stolz, M.S. Sacks, B.B. Keller, W.R. Wagner, An elastic, biodegradable cardiac patch induces contractile smooth muscle and improves cardiac remodeling and function in subacute myocardial infarction, J. Am. Coll. Cardiol. 49(23) (2007) 2292–300.

[19] J. Ando, K. Yamamoto, Flow detection and calcium signalling in vascular endothelial cells, Cardiovasc. Res. 99(2) (2013) 260–8.

[20] S. Pok, I.V. Stupin, C. Tsao, R.G. Pautler, Y. Gao, R.M. Nieto, Z.-W. Tao, C.D. Fraser Jr., A.V. Annapragada, J.G. Jacot, Full-Thickness Heart Repair with an Engineered Multilayered Myocardial Patch in Rat Model, Advanced Healthcare Materials 6(5) (2017) 1600549.

[21] Q. Liu, S. Tian, C. Zhao, X. Chen, I. Lei, Z. Wang, P.X. Ma, Porous nanofibrous poly(l-lactic acid) scaffolds supporting cardiovascular progenitor cells for cardiac tissue engineering, Acta Biomater. 26 (2015) 105–114.

[22] R. Hashizume, Y. Hong, K. Takanari, K.L. Fujimoto, K. Tobita, W.R. Wagner, The effect of polymer degradation time on functional outcomes of temporary elastic patch support in ischemic cardiomyopathy, Biomaterials 34(30) (2013) 7353–63.

[23] J. Xie, Y. Yao, S. Wang, L. Fan, J. Ding, Y. Gao, S. Li, L. Shen, Y. Zhu, C. Gao, Alleviating Oxidative Injury of Myocardial Infarction by a Fibrous Polyurethane Patch with Condensed ROS-Scavenging Backbone Units, Adv Healthc Mater 11(4) (2022) e2101855.

[24] Y. Yao, J. Ding, Z. Wang, H. Zhang, J. Xie, Y. Wang, L. Hong, Z. Mao, J. Gao, C. Gao, ROS-responsive polyurethane fibrous patches loaded with methylprednisolone (MP) for restoring structures and functions of infarcted myocardium in vivo, Biomaterials 232 (2020) 119726.

[25] A. D’Amore, T. Yoshizumi, S.K. Luketich, M.T. Wolf, X. Gu, M. Cammarata, R. Hoff, S.F. Badylak, W.R. Wagner, Bi-layered polyurethane - Extracellular matrix cardiac patch improves ischemic ventricular wall remodeling in a rat model, Biomaterials 107 (2016) 1–14.

[26] M. Shafiq, Y. Zhang, D. Zhu, Z. Zhao, D.H. Kim, S.H. Kim, D. Kong, In situ cardiac regeneration by using neuropeptide substance P and IGF-1C peptide eluting heart patches, Regenerative biomaterials 5(5) (2018) 303–316.

[27] Y. Liu, Y. Xu, Z. Wang, D. Wen, W. Zhang, S. Schmull, H. Li, Y. Chen, S. Xue, Electrospun nanofibrous sheets of collagen/elastin/polycaprolactone improve cardiac repair after myocardial infarction, American journal of translational research 8(4) (2016) 1678–94.

[28] R. Hashizume, K.L. Fujimoto, Y. Hong, J. Guan, C. Toma, K. Tobita, W.R. Wagner, Biodegradable elastic patch plasty ameliorates left ventricular adverse remodeling after ischemia-reperfusion injury: a preclinical study of a porous polyurethane material in a porcine model, J. Thorac. Cardiovasc. Surg. 146(2) (2013) 391–9 e1.

[29] L.M. Silveira-Filho, G.N. Coyan, A. Adamo, S.K. Luketich, G. Menallo, A. D’Amore, W.R. Wagner, Can a Biohybrid Patch Salvage Ventricular Function at a Late Time Point in the Post-Infarction Remodeling Process?, JACC: Basic to Translational Science 6(5) (2021) 447–463.

[30] C.X. Lam, M.M. Savalani, S.H. Teoh, D.W. Hutmacher, Dynamics of in vitro polymer degradation of polycaprolactone-based scaffolds: accelerated versus simulated physiological conditions, Biomed. Mater. 3(3) (2008) 034108.

