## supplemental for "Cardiac patch treatment alleviates ischemic cardiomyopathy correlated with reverting Piezo1/2 expression by unloading left ventricular myocardium"

| Table 1. |  |
| --- | --- |
| Primer | 5' to 3' |
| *Atp1a1*-forward | ACCTGCCTATCCTTAAGCGTG |
| *Atp1a1*-reverse | CCGATGCGTTTGGGTTCTTG |
| *Angpt2*-forward | AGAGTACAAAGAGGGCTTCGG |
| *Angpt2*-reverse | TCCTGGTTGGCTGATGCTAC |
| *Piezo1*-forward | TACTGGCTGTTGCTACCCTG |
| *Piezo1*-reverse | CACGTTTGCCCAAAGGTTACAG |
| *Piezo2*-forward | CAACCAAAGCGACGATGCAA |
| *Piezo2*-reverse | CCAGCATCAGCTCCCTTCAA |
| *18s*-forward | ACCGCAGCTAGGAATAATGGA |
| *18s*-reverse | GCCTCAGTTCCGAAAACCA |

**For quantitative real time RT-PCR in rat, the following primers were used:**
